# Agonists to ions: Efficiency suggests ‘zipper’ mechanism for nicotinic receptor activation

**DOI:** 10.1101/2021.12.18.473297

**Authors:** Anthony Auerbach

## Abstract

Agonists are classified by the strength at which they bind to their target sites (affinity) and their ability to activate receptors once bound to those sites (efficacy). Efficiency is a third fundamental agonist property that is a measure of the correlation between affinity and efficacy. Efficiency is the percent of agonist binding energy that is converted into energy for receptor activation (‘gating’). In the muscle nicotinic acetylcholine receptor, agonists belong to families having discrete efficiencies of 54%, 51%, 42% or 35%. Efficiency depends on the size and composition of both the agonist and binding site, and can be estimated from, and used to interpret, concentration-response curves. A correlation between affinity and efficacy indicates that the agonist’s energy changes that take place within binding and gating processes are linked. Efficiency suggests that receptors turn on and off by progressing through a sequence of energy-linked domain rearrangements, as in a zipper.

---

Nicotinic acetylcholine receptors (AChRs) have long served as a model system for understanding how drugs activate receptors. Many essential concepts in pharmacology (receptor, agonist, affinity, efficacy, desensitization) originated from studies of AChRs. Below, I review fundamental aspects of receptor operation and show how 3 unexpected experimental results (Figs. 3, 6 and 8) illuminate agonist efficiency and a ‘zipper’ mechanism for AChR gating.

**Figure 1.**
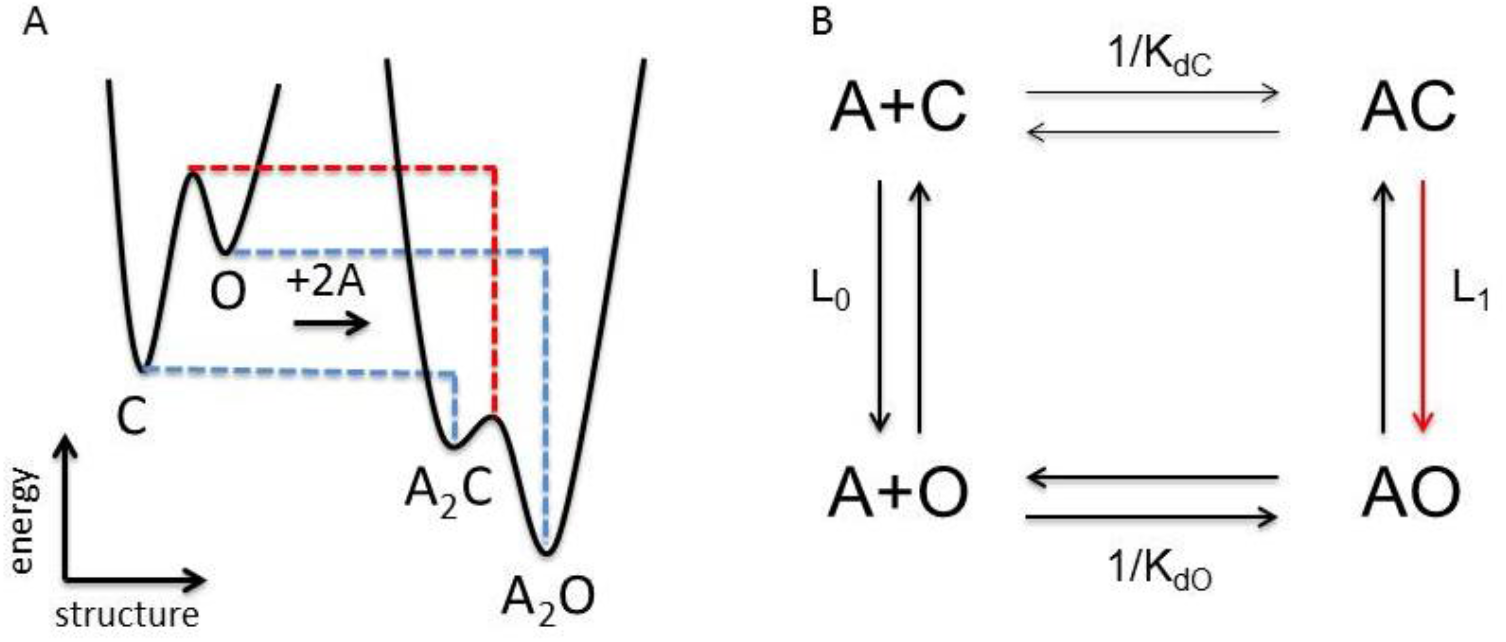
Agonists stabilize the gating transition state. A. Potential energy surfaces (landscapes) for gating. Resting (C) and active (O) shapes are shown as energy wells separated by an energy barrier (the TS, red line). C has a closed channel and low-affinity binding sites. O has an open channel and high-affinity binding sites. Left, without bound agonists (A) the TS and O energies are high (relative to C) and the opening rate constant and probability of being O (P_O_) are small. Right, agonist binding energy (dashed lines) preferentially stabilizes the TS and O, to increase the opening rate constant and P_O_. B. Cyclic reaction scheme for a receptor with 1 agonist binding site. L_0_ and L_1_, C⟷O gating equilibrium constant without and with a bound agonist; K_dC_ and K_dO_, equilibrium dissociation constant of the C and O conformation. The product of equilibrium constants that connect diagonal states are equal: L_1_/K_dC_=L_0_/K_dO_ (see Eq. 1). Red arrow, a faster opening rate constat (from stronger agonist binding to the TS) is the reason L_1_>L_0_.

**Figure 2.**
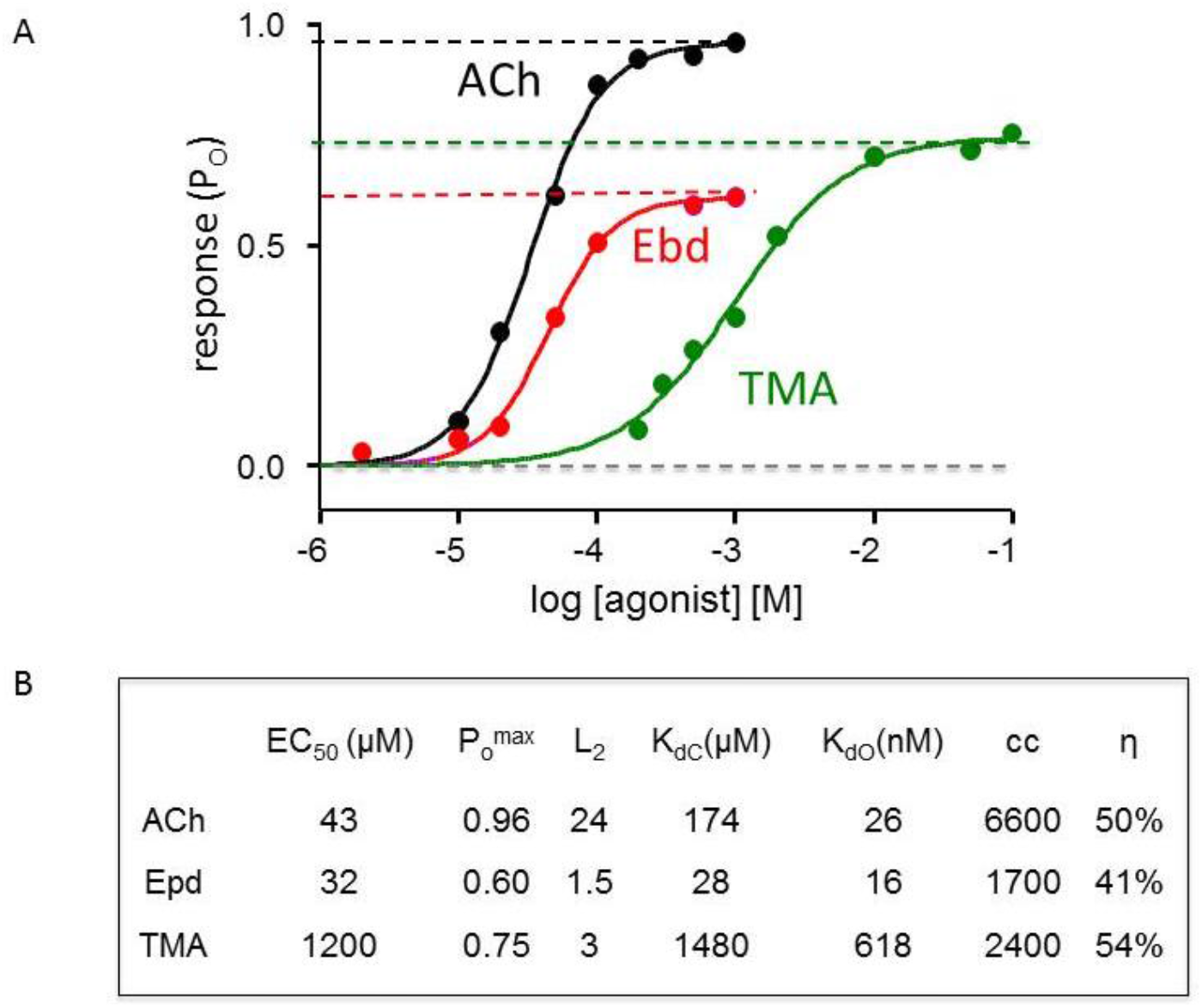
Estimating efficiency from a CRC. A. Single-channel open probability P_O_ measured at different agonist concentrations and fitted by the Hill equation to estimate a high concentration asymptote (max; dashed lines) and EC_50_ (concentration where P_O_ is half-max). The low concentration asymptote (min; gray dashed line) is the same for all agonists and too small to measure by the fit. ACh, acetylcholine; Ebd, epibatidine; TMA, tetramethylammonium (muscle adult-type AChRs, -100 mV). B. Table of fitted CRC parameters and calculated equilibrium constants (Eq. 2; L_O_ equal to 7 × 10^−7^). cc, coupling constant (=K_dC_/K_dO_); η, efficiency (=1-logK_dC_/logK_dO_).

**Figure 3.**
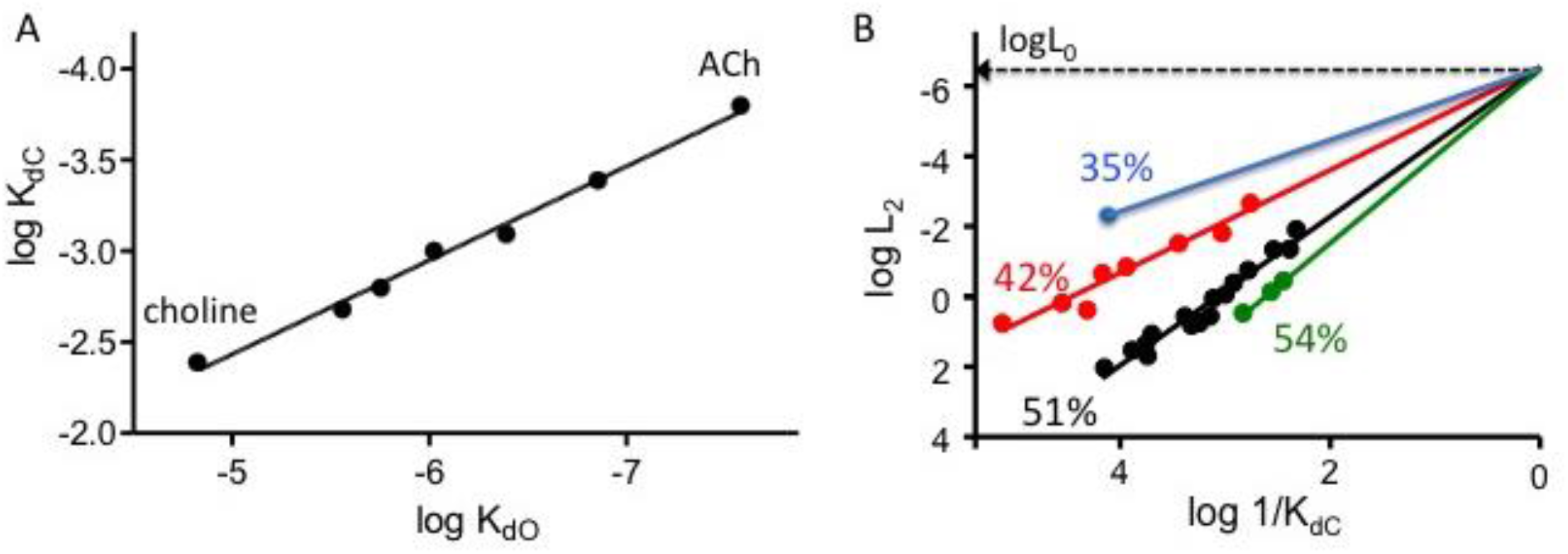
Efficiency. A. Agonist binding to O is twice as strong as to C regardless of agonist affinity, potency or efficacy (Jadey & Auerbach, 2012). (Binding energy is proportional to the logarithm of an equilibrium association constant.) All of the agonists fall on the same line, slope K_dC_/K_dO_=0.51 ± 0.02, indicating that all have the same efficiency (Eq. 3). B. Efficiency plot. The x-axis is proportional agonist binding energy (log affinity) and the y-axis is proportional to the coupling constant energy (log efficacy)(Eq. 1). The red, black and green lines were fitted independently by Eq. 4 to yield slopes that estimate efficiency and y-intercepts that pertain to L_0_. Affinity and efficacy are correlated on a log-log scale, but only for agonists that have the same efficiency. The difference between 54% and 50% populations is significant (Fig. 5). Agonists: green - TMA, dimethylpyrrolidinium, dimethythiazolidinium; black - ACh, carbamylcholine, choline, 3-hydroxyproplyTMA, 4-hydroxybutylTMA, nicotine, nornicotine, dimethyhlphenylpiperizinium (DMPP), anabaesine; red - epibatidine, epiboxidine, cytisine, azabicycloheptane anatoxin-A, tetraethylammonium, tetramethylphosphonium. The efficiency of varenicline (blue) was calculated using L_0_ estimated from the other y-intercepts.

**Figure 4.**
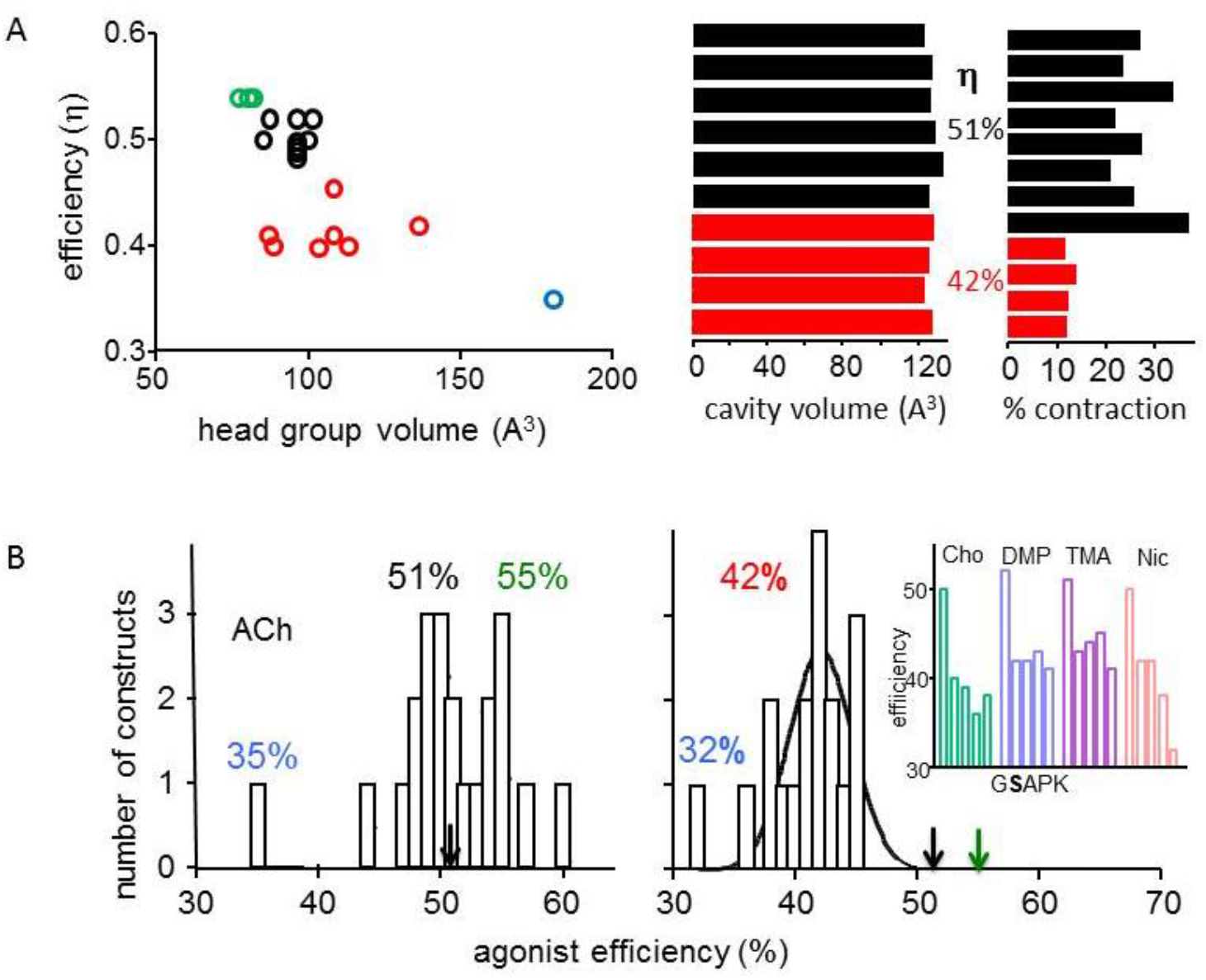
Structural correlates of efficiency. A. In AChRs, efficiency is correlated inversely with agonist head-group volume (left) and depends on the extent of contraction of the binding cavity in C→O (right) (color code, Fig. 3). B. Mutations of residues in the adult-type binding cavity influence efficiency (arrows mark wt values). Left, most mutations of 4 aromatic residues in the *α* subunit have no effect on ACh efficiency, but some increase (W149A, rightmost) and one decreases (Y190A, leftmost) significantly. Right, mutations of a conserved glycine near the binding cavity (*α*G153) reduce efficiency to 42%. Inset, agonist/mutation combinations; S (bold) causes a congenital myaesthenic syndrome (Sine et al., 1995).

**Figure 5.**
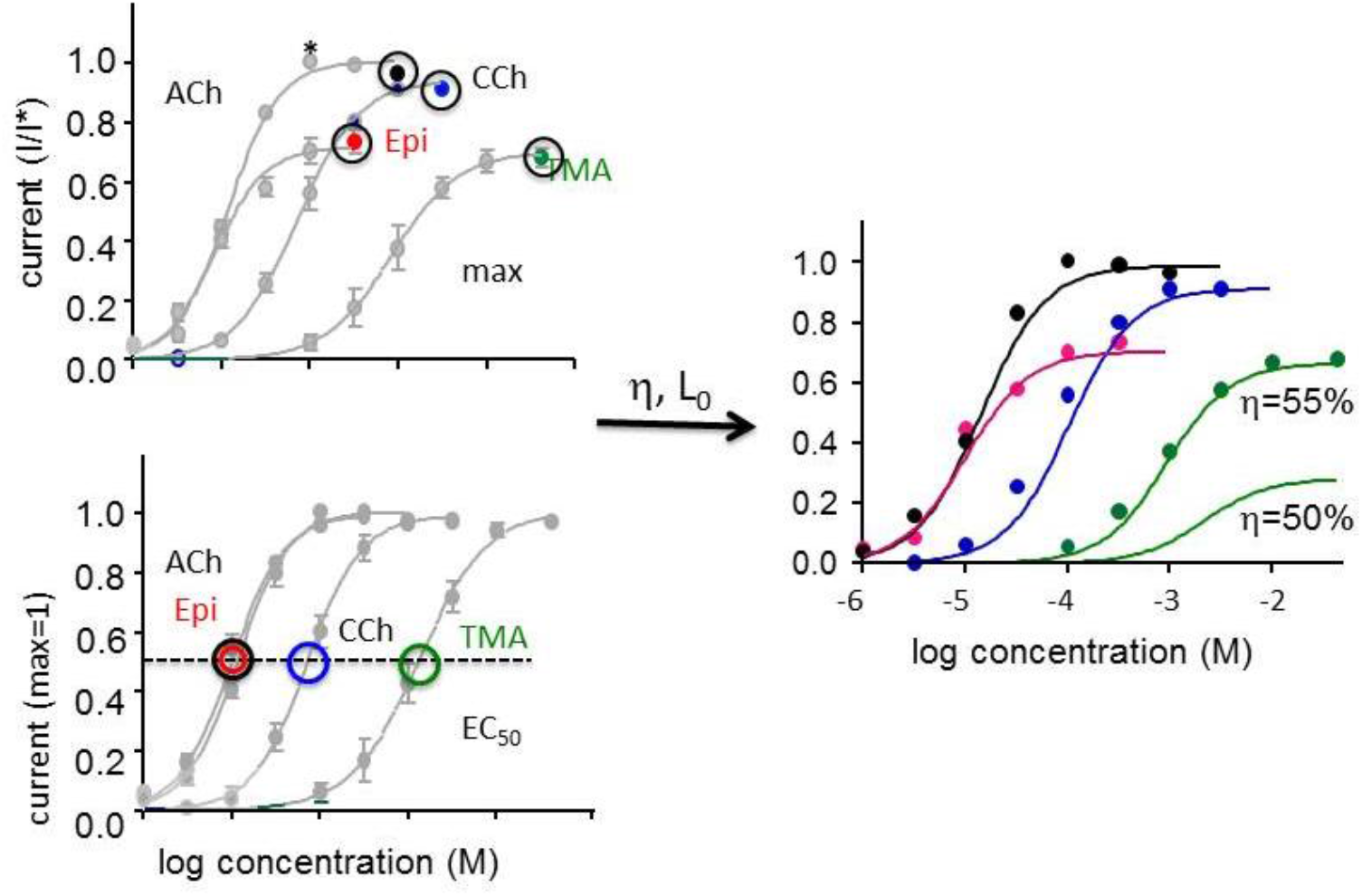
Efficiency applied to CRC analysis. Left, CRCs constructed from whole-cell current responses (adult-type AChRs) (ref). Top, responses normalized to that at 300 μM ACh (*); max values are circled. Bottom, responses normalized to max=1; EC_50_ values are circled. Right, CRCs (solid curves) calculated from η (42 or 50%), L_0_ (7× 10^−7^) and either max or EC_50_ (left panels). The calculated curve for TMA matches when η is 55%.

**Figure 6.**
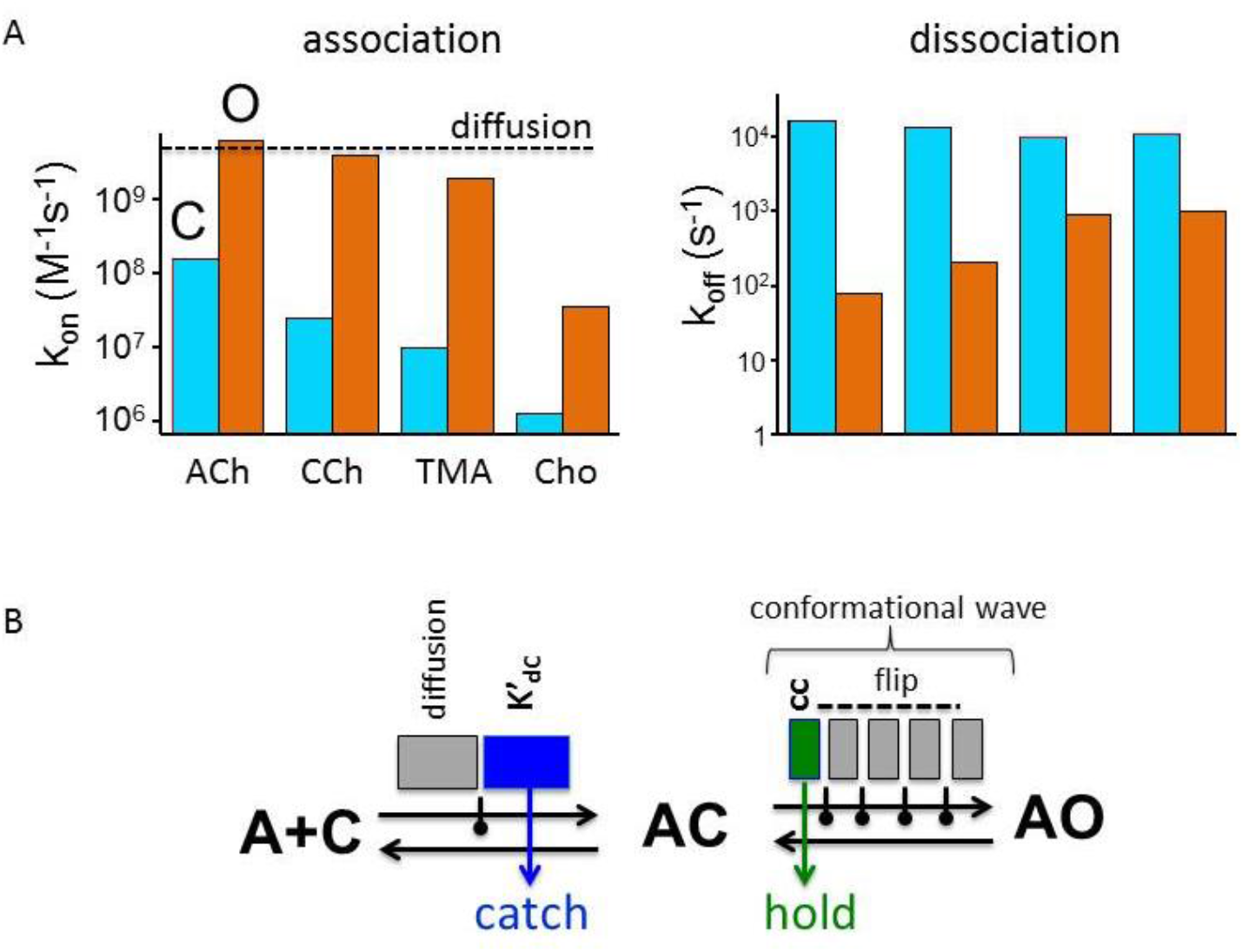
Intermediate transitions in receptor activation. A. Agonist binding rate constants (Nayak & Auerbach, 2017). Left, association (k_on_) to O (orange) is near the diffusion limit whereas k_on_ to C (blue) is slower and agonist-dependent. Right, dissociation (k_off_) from C is approximately the same for all the agonists. k_off_ from O is slower and more agonist-dependent. The agonist-dependent component of K_dC_ (affinity) derives from differences in k_on_. B. There are intermediate transitions and states in both binding and gating. Left, binding. K_dC_ is a function of both diffusion (gray bar) and a local conformational change at the binding site (‘catch’; blue bar). The equilibrium constant for this rearrangement (K’_dC_) determines relative affinity. Right, gating. There are 5 intermediate transitions in the isomerization that comprise a conformational wave (see Fig. 8). In opening only the first is agonist dependent (‘hold’, green bar) in which there is an increase in binding energy. The equilibrium constant for this rearrangement (cc) determines relative efficacy. The remaining intermediate transitions (gray bars) are rearrangements away from the binding site and are not agonist dependent. Only the agonist energy changes in catch and hold (blue and green bars) determine efficiency. Black dots, intermediate states connected by the transitions. Flip possibly reflects a sojourn in one or more gating intermediate.

**Figure 7.**
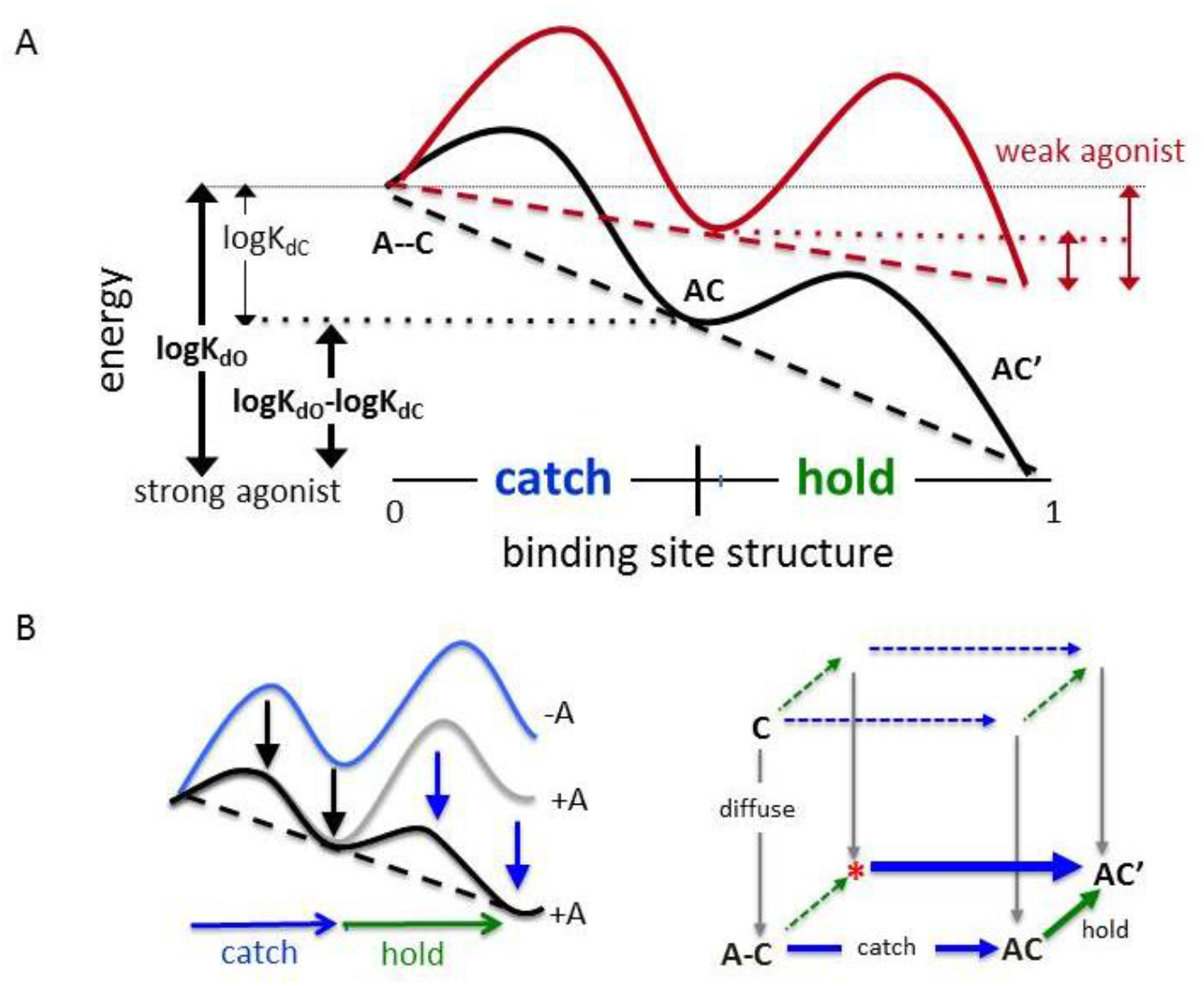
Energy landscapes and models of agonist binding. A. Potential energy surfaces for a strong (black) and a weak agonist. The landscapes pertain only to catch and hold, the intermediate transitions in binding and gating (Fig. 6B). Left vertical arrows show agonist energy changes. η, the fraction of the total energy change applied to hold, is the same for both strong and weak agonists. Agonists with different affinities and efficacies have the same efficiency because catch and hold are linked in a linear free energy relationship (LFER; dashed tilted line). B. Catch and hold promote each other because of the LFER. Left, The presence of an agonist lowers the energy of the catch barrier and state (black arrows), which in turn lowers the energy of the hold barrier and state (blue arrows). Right, the primary activation pathway is diffuse-catch-hold; arrow thickness indicates equilibrium constant value. The structure hold-without-catch is rare, but microscopic reversibility (bottom cycle) dictates that catch will occurs with a high probability.

**Figure 8.**
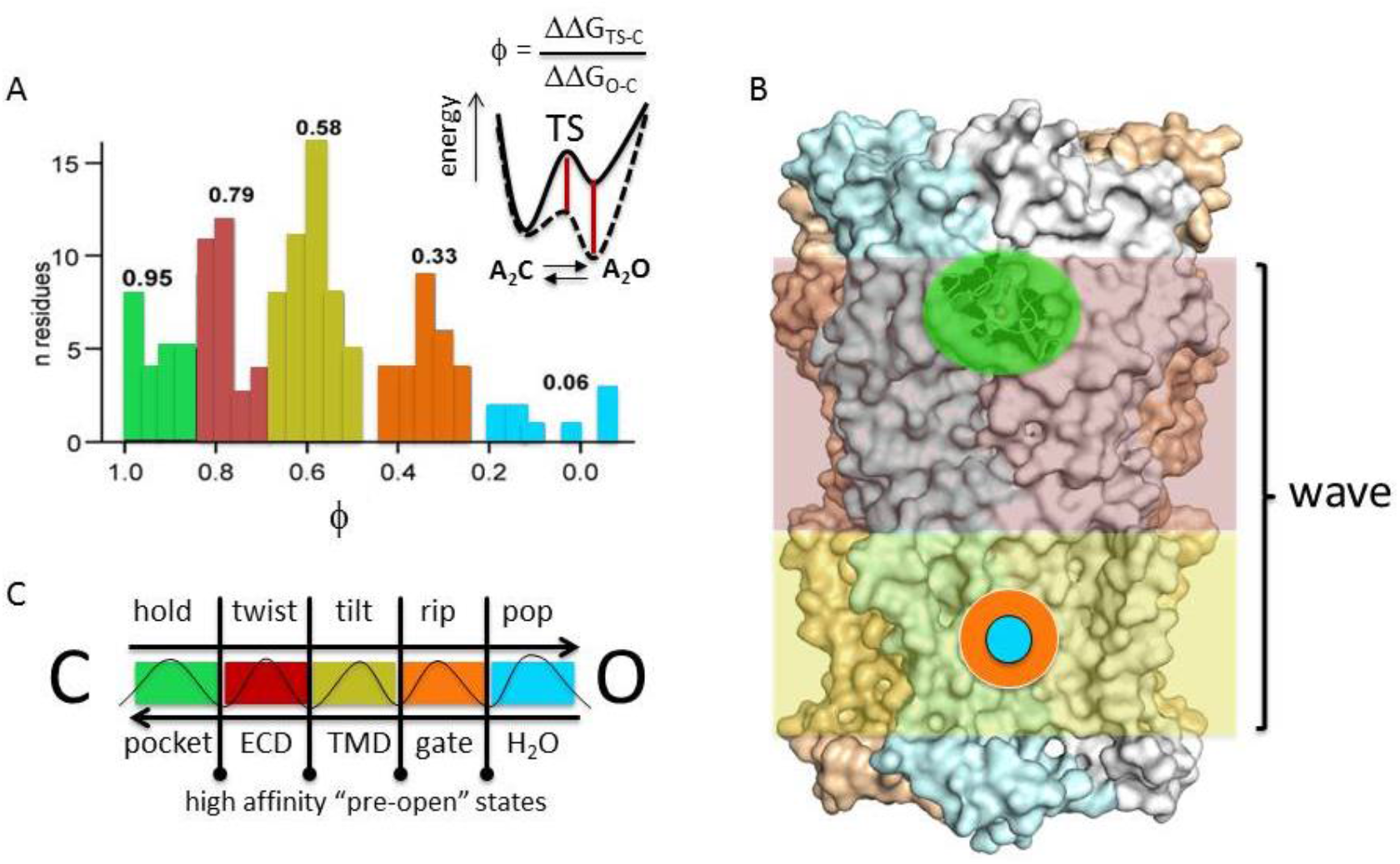
The gating transition state. Energy at the gating TS is organized with regard to distribution and map (Purohit et al., 2013). A. ϕ is the TS/O energy ratio (inset, red lines) estimate for a series of mutations of one amino acid. There are 5 distinct populations of residue ϕ values. B. Map of ϕ values. Each ϕ population is associated with a different domain of the *α* subunit: green, binding pocket; red, extracellular domain (ECD); yellow, transmembrane domain (TMD); orange, gate region; blue, water. C. The ϕ populations correspond to passage across intermediate transitions that together comprise a gating conformational ‘wave’ (Grosman, Zhou, & Auerbach, 2000). There are 4 high-affinity intermediate states (lines with dots) that separate 5 intermediate transitions (names above and locations below) that reflect local structural rearrangements of 5 protein domains.

**Figure 9.**
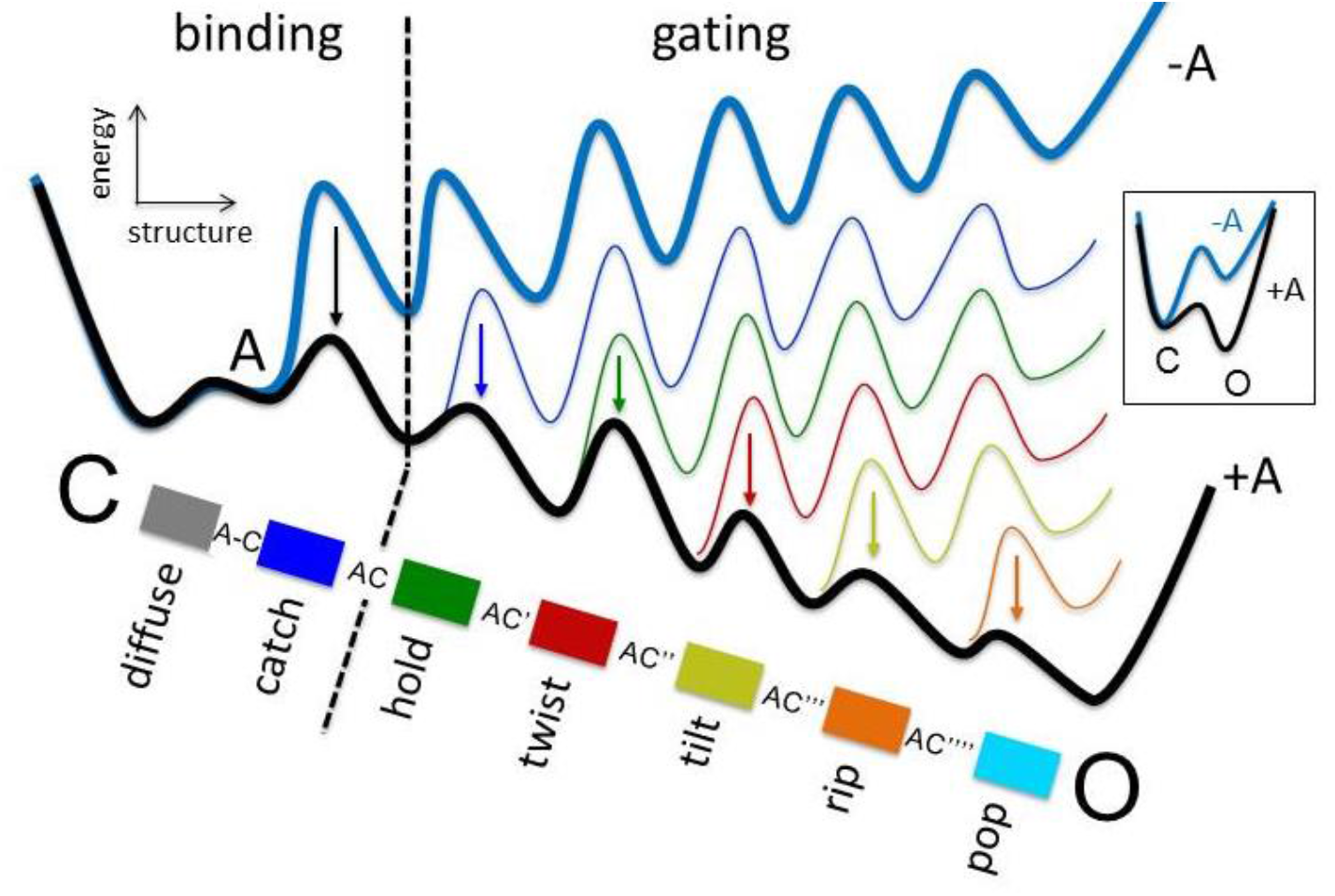
Energy landscape for the binding-gating zipper. As in a zipper, each transition in the gating conformational wave lowers the barrier for the next, in sequence (arrows) (see Fig. 7B, left). The top (blue) landscape, no agonist. The series of intermediate rearrangements in opening, catch-hold-twist-tilt-rip-pop, are linked by a series of LFERs. The transitions occur in the reverse order in channel closing because of microscopic reversibility. As a consequence, increasing constitutive P_O_ by adding mutations enforces the reversal of all previous elements in the chain including catch (crossing the binding-gating divide; dashed line), so that k_on_ to O becomes limited by diffusion (Fig. 6A). The sequential protein rearrangements that connect agonist diffusion and water fusion include both binding and gating events and are linked structurally and energetically to generate a single conformational sweep. Bottom, there are 4 intermediate gating states all of which have a high affinity for the agonist. Inset, at low time resolution the intermediate transitions merge into a single barrier.

## 1. Efficiency: definition, measurement, applications

### Receptor activation by agonists

Allosteric proteins alternate between global conformations that have different functional outputs and different sensitivities to environmental stimuli (Changeux & Edelstein, 1998; Nussinov & Tsai, 2013). Ligand-gated ion channels alternate between shapes called C and O, that have a pore that is closed or open and ligand sites that bind agonists weakly or strongly. Stronger binding to O serves to increase the probability that the receptor adopts the open-channel conformation (P_O_). Agonists activate receptors because they contribute binding energy that preferentially stabilizes O as well as intermediates structures that populate the C⟷O conformational-change pathway.

This scenario for receptor activation can be depicted either as potential energy surfaces or as a cyclic reaction scheme (Fig. 1). In the energy landscapes, C and O are shown as energy wells (regions of stability) separated by an energy barrier (a region of instability) called the transition state (TS). Without bound agonists, the TS barrier is large and O is unstable, so both the opening rate constant and P_O_ are small. The P_O_ of unliganded adult muscle nicotinic acetylcholine receptors (AChRs) is ∼10_-6_ with openings occurring once every ∼15 minutes. From a distance, it looks like there is no activity whatsoever.

The situation changes when agonists are bound to the C conformation. Both the TS and O are stabilized almost equally by the extra binding energy, to flatten the barrier and facilitate a spontaneous transition from C to O. The reverse, channel-closing process is hardly affected because the difference between the TS and O energies remains essentially the same. In AChRs, after 2 neurotransmitter (ACh) molecules have bound with low affinity to C, the switch of the binding sites from low to high affinity at the onset of the global isomerization causes P_O_ to rise to ∼0.96 and opening transitions to occur within ∼20 μs. Although agonist binding energy does stabilize O slightly more than the TS, the ∼4-fold prolongation of the open-channel lifetime is insignificant compared to the ∼40 million-fold increase in the channel-opening rate constant.

These energy landscapes can be displayed as a cyclic reaction scheme in which vertical steps represent the C⟷O conformational change (‘gating’) and horizontal steps represent formation of an agonist-protein complex (‘binding’). More generally, vertical steps represent the global change in protein shape (and function) and horizontal steps represent the deposition of environmental energy (a stimulus) into the alternative shapes. Evidence presented below shows that binding and gating are not separate enterprises but rather are different stages of a single, energy-linked process.

It is usually assumed that all transitions in the cycle (movements along the potential energy surfaces) occur spontaneously, driven only by temperature (Frauenfelder, 2014). An agonist does not activate a receptor by transferring momentum to the protein. Rather, a diffusing ligand is in thermal equilibrium and enters its binding pocket gently and randomly. Accordingly, the sum of energy changes (product of equilibrium constants) is the same for clockwise and counter clockwise pathways connecting the corners of the cycle. While it is possible that in some receptors external energy sources other than temperature are present, in AChRs microscopic reversibility has been demonstrated experimentally so these are insignificant (Nayak & Auerbach, 2017).

Unliganded C receptors constantly undergo thermal motions but rarely succeed in surmounting the high gating TS to reach O. After the agonist has entered the binding pocket and formed a low-affinity complex, the system jitters randomly towards the TS thereby shrinking the separating barrier as favorable binding energy is added. That is, agonists bind more strongly to both TS and O structures, compared to C. With agonists in the binding sites, the pathway from C to O is leveled so that a spontaneous transition to the open-channel conformation can occur rapidly. Although gating TS structures of receptors have not yet been solved, it is possible that, as with enzymes, a favorable match between ligand and protein structures at the TS stabilizes the system, easing the spontaneous transition to O.

### CRCs

Fig. 2A shows dose (concentration)-response curves (CRCs) for 3 AChR agonists. The responses were fitted by an S-shaped empirical function to estimate a high-concentration asymptote (max) and midpoint (EC_50_). In pharmacology terms, max reflects agonist efficacy and EC_50_ is the inverse of agonist potency that in turn relates to agonist affinity (weak binding to C). An additional CRC parameter is the low-concentration asymptote (min) that is the same for all agonists and usually too small to be measured directly from a fit. We will make use of this parameter, with its value estimated *a priori* using different methods. Another CRC parameter that is estimated the fit is the slope of the curve at the midpoint (Utkin et al.) that is proportional to the number of agonist binding sites (n) (Qin, 2004) and is also known *a priori*.

The CRC parameters (and pharmacology terms) relate to equilibrium constants of the cycle (Cechini). In adult-type (mouse and human) AChRs there are 2 binding sites that have similar affinities for the agonists we will consider here (cyclic). Because microscopic reversibility is satisfied,

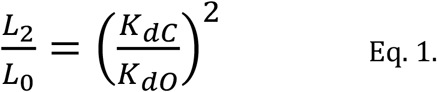

L is a gating equilibrium constant (subscript indicates number of bound agonists) and K_d_ is an agonist equilibrium dissociation constant (subscript indicates receptor conformation). Although there are 2 agonist “affinities”, 1/K_dC_ and 1/K_dO_, the word refers just to the lower affinity, 1/K_dC_. The high/low affinity ratio K_dC_/K_dO_ is called the ‘coupling’ constant (cc) and is the factor by which each agonist molecule increases the gating equilibrium constant.

When the agonist increase in P_O_ depends mainly on binding to C and gating with bound agonists, the following equations relate CRC parameters to equilibrium constants of the cycle (2 equivalent binding sites) (Cecchini & Changeux, 2021; Indurthi & Auerbach, 2021):

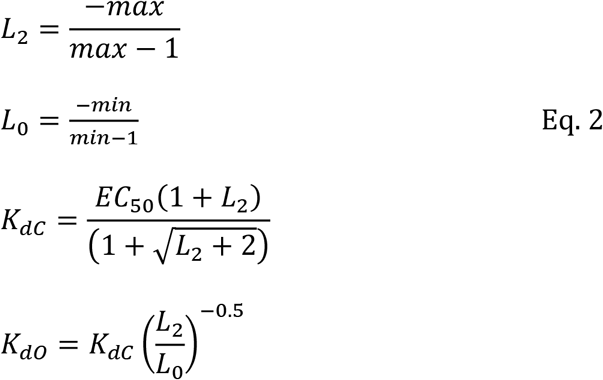

Hence, all 4 equilibrium constants of the cycle can be calculated from the high-concentration asymptote (efficacy), zero-concentration asymptote (unliganded activity) and midpoint (potency) of a CRC.

In practice, receptors also adopt conformations that are not included in the cycle. For instance, they can be inactive but have a high affinity for agonists (desensitized) or have the O shape but not generate a response (blocked). Certainly, sojourns in such extra-cycle states influence P_O_ and the cellular CRC. However, our goal here is to understand the chemical mechanisms that undergird receptor activation rather than to describe physiological responses, so I will focus on only the binding-gating events described by the cycle. Accordingly, in results described below extra-cycle events have been largely excluded from the experimental measurements to allow a clear accounting of just agonist binding and receptor gating.

The fitted parameters and calculated equilibrium constants for the 3 CRCs are given in Fig. 2B. The value of the unliganded gating constant (L_0_, related to min) measured previously (Nayak, Purohit, & Auerbach, 2012; Purohit & Auerbach, 2009) is tiny (∼7 × 10^−7^ at -100 mV in adult-type AChRs) and was not obtained from the CRC. For all three agonists, K_dC_ (weak binding) is μM and K_dO_ (strong binding) is nM. L_0_ and n are the same for all ligands, so the cc alone determines the agonist-dependent component of L_2_ and, therefore, relative agonist efficacy. Note that is necessary to know L_0_ in order to calculate the cc.

Coupling constants have been measured for many AChR agonists and fall in the range ∼5 to ∼6000 (Bruhova; Jadey). The cc for choline, an ACh breakdown product that is present at high concentrations at cholinergic synapses, is ∼250. Whereas 2 ACh bound molecules increase L_2_ over the baseline by a factor of ∼36 million (P_O_ max ∼0.96), 2 bound choline molecules increase L_2_ only by a factor of ∼60 thousand (P_O_ max ∼0.05).

An agonist is a molecule with cc>1 (K_dC_>K_dO_). These ligands increase the gating equilibrium constant and generate a response above the baseline (Eq. 1). An antagonist is a molecule with cc=1 (K_dC_=K_dO_). These ligands bind competitively but neither increase or decrease activity. An inverse agonist is a molecule with cc<1 (K_dC_<K_dO_). These ligands reduce the gating equilibrium constant and throttle receptor activity.

In some receptors there is a brief shut state (∼μs lifetime) that, in the cycle, could be interposed between C and O (Auerbach, 1993; Dmitrij Ljaschenko, 2021; Lape, Colquhoun, & Sivilotti, 2008; Mukhtasimova, Lee, Wang, & Sine, 2009). Indeed, from what we know about conformational changes in large multimeric proteins many such short-lived states must exist with < μs lifetimes, too brief to be resolved directly by using electrophysiology. These intermediate transitions and state can be represented by adding small hills and valleys (corrugations) to the potential energy surface of the C⟷O isomerization (Fig. 1A). However, the agonist’s cc is unaltered by transit through these because the K_dC_/K_dO_ ratio depends only on the relative free energies of the gating end states, O versus C (Eq. 1). When the cycle applies, the relative efficacy of an agonist is independent of reaction intermediates.

A CRC can be thought of as a plot of energy-in (related to agonist concentration) versus energy-out (related to the response). From Fig. 2A, the neurotransmitter ACh and the frog toxin Ebd have a similar energy-in (potency) but ACh produces a significantly greater energy-out (max). Thus, by inspection alone we predict that a greater fraction of ACh’s binding energy is converted into gating energy. This fraction defines agonist efficiency (η; eta). TMA has both a lower potency and max compared to ACh so its relative η cannot be determined by inspection. To do this, we need to delve deeper into efficiency.

### Efficiency

The first unexpected result is shown in Fig. 3A. In muscle AChRs, the strong/weak binding energy ratio K_dC_/K_dO_ is the same for a series of agonists regardless of affinity or efficacy. The thermodynamic cycle dictates only that for agonist to activate the receptor it must bind more strongly to O versus C but says nothing more about the relationship between binding energies. Therefore, it was a surprise to learn that for both partial and full agonists, binding energy approximately doubles upon receptor activation.

In thermodynamics, the efficiency of a machine is defined as the energy-out/energy-in ratio. For simplicity, we will first consider a receptor that has only 1 agonist binding-site. From Eq. 1, the useful energy-out (activity above the baseline) is proportional to minus log of the coupling constant, or (logK_dO_-logK_dC_). The maximum energy-in is from binding to the higher-affinity conformation and is proportional to (logK_dO_). Taking the ratio (Nayak, Vij, Bruhova, Shandilya, & Auerbach, 2019),

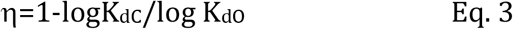

The results in Fig. 3A show that the agonists all have the same efficiency. The efficiency of a steam engine has the same form as Eq. 3 but with the hot/cold temperature ratio replacing the strong/weak binding energy ratio. Note that agonist efficacy and efficiency are distinct, with the former being a ratio of binding constants and the latter being a ratio of binding energies. Plugging K_d_ values into Eq. 3 and multiplying by 100% gives efficiency as a percentage.

From the CRC parameters (Fig. 2), the agonist η-values are 50% for ACh, 41% for Ebd and 54% for TMA. As predicted by inspection of the CRC, the neurotransmitter applies a greater percentage of its binding energy to gating compared to the frog toxin. Eq. 3 allows us to estimate that ACh is 8% more efficient than Epi and that TMA is 4% more efficient than ACh.

The efficiency of any agonist at any receptor binding-site can be calculated by using Eq. 3 once the two binding constants are known. As described above (Indurthi & Auerbach, 2021), these can be estimated from fitting a CRC if L_0_ is known *a priori*. There are, however, several alternative approaches that do not require prior knowledge of L_0_. K_dC_ and K_dO_ can be estimated directly from single-channel kinetic currents (Nayak & Auerbach, 2017) or, possibly, from *in vitro* binding assays. In the following paragraphs I show that in addition η can be estimated without knowledge of L_0_ from CRCs for a group of agonists.

Although it is interesting and possibly useful to quantify the fraction of binding energy each ligand applies to receptor activation, η is much more valuable because agonists having different affinities and efficacies can have the same efficiency (Jadey & Auerbach, 2012). For instance, in AChRs both the partial agonist choline (max=0.05, EC_50_=4 mM) and the full agonist ACh (max=0.96; EC_50_=170 μM) have an efficiency of 50%. ACh and choline both apply half of their binding energy to gating, even though ACh has much more to give. Ebd and its congers (see below) apply only 42% of their binding energy to receptor activation. Efficiency is not only distinct from affinity and efficacy, it also is a new way to classify agonists.

To better delineate efficiency classes, a log-log plot of binding versus gating constant is useful (Nayak et al., 2019). Combining Eqs.1 and 2,

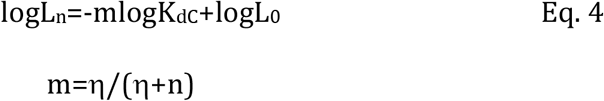

which is an equation for a straight line, y=mx+b, with n the number of (equivalent) binding sites. An ‘efficiency’ plot, logL_2_ versus log(1/K_dC_) for a group of agonists that have the same efficiency, is a series of points that fall on the same line, the slope of which depends on efficiency. The x and y-axes of this plot are binding versus gating energies, so each point represents the correlation between affinity and efficacy for that agonist and reflects an entire CRC.

Fig. 3 is an efficiency plot for 20 AChR agonists. All but one fall on a line having one of 3 slopes that correspond to average η values that are the same as those in Fig. 2B (54%, 51% or 42%). There are several important features of this plot to consider.

First, efficiency classes are discrete. With one exception, agonist efficiencies are not scattered but rather align tightly to one of the three classes. The 54% and 51% classes are different (see Fig. 5), so efficiency values are also precise.

Second, although the points from each efficiency class were fitted independently the 3 lines converge to the same y-intercept that is equal to log L_0_ (Eq. 4). As expected, the unliganded gating equilibrium constant is agonist-independent. Moreover, L_0_ estimated from the efficiency plot is the same as the values estimated previously using completely different methods (Purohit, Nayak, Jha). Hence, the efficiency plot is a way to estimate L_0_ from readily-measured CRC parameters, max and EC_50_.

A third important feature of Fig. 4 is that the correlation (in energy) between affinity and efficacy is apparent only if one considers agonist efficiency. If every agonist had a unique efficiency the points would be scattered and concept of agonist efficiency would be less useful. And, if only a handful of agonists (each from an entire CRC) were plotted, the correlations would probably not be apparent. The correlation and segregation into η classes is clear only when many agonists are plotted.

Based on literature reports of affinity and efficacy, these properties appear to be correlated in agonists other receptors (glutamate, glycine, GABA_A_ and muscarinic) and so generate linear efficiency plots (citations given in (Nayak et al., 2019)). The efficiencies (49%, 59%, 39% and 52%, respectively) and L_0_ values (2×10^−3^, 5×10^−4^, 4×10^−2^, and 1×10^−4^) estimated from the y-intercepts of these efficiency plots are reasonable. It is possible that the consensus that there is no correlation between affinity and efficacy can be traced to examining only a few agonists and not considering that agonists belong to different efficiency classes. Scatter in a log-log plot of affinity vs. efficacy does not demonstrate a lack of correlation, but rather only that agonists can belong to families that have different correlations.

In Fig. 3, one AChR agonist, varenicline, did not fall on any of the 3 lines. Using the L_0_ value obtained from the other agonists we calculate its efficiency to be 35%. It is possible other agonists have this efficiency. A mutations in the vicinity of the transmitter binding site lowers the efficiency of ACh to around the level (see below).

From Eq. 3, if an agonist’s efficiency is known, K_dC_ and K_dO_ can be calculated from each other. This means that if the y-intercept of the efficiency plot (L_0_, the same for all agonists) is also known, EC_50_ and max can be calculated from each other (Eq. 2). Accordingly, given η and L_0_, an entire CRC can be calculated from just the maximum response, and efficacy information can be regained from a CRC that has been normalized to have a maximum response of 1 (Fig. 4). The requirement that L_0_ and η must be known may not be overly burdensome. L_0_ has to be measured only once for each receptor (perhaps from an efficiency plot), and η is the same for a group of ligands and possibly can be inferred from agonist structure (see below).

To summarize, η is the fraction of energy from agonist binding that is converted into energy for receptor gating. Efficiency is a new way to categorize agonists that are now classified only by affinity and efficacy. η allows an estimate of L_0_ from easy-to-measure CRC parameters, and an entire un-normalized CRC to be calculated from either the response to just 1 agonist concentration (max) or EC_50_ of a normalized response. In the next sections I describe structural correlates of efficiency in AChRs, and discuss a deep implication of the affinity-efficacy correlation that illuminates the receptor activation mechanism.

## 2. Efficiency: structural correlates and a zipper mechanism for gating

### Structural correlates of efficiency

There is an inverse relationship between the volume of the agonist’s head group and its efficiency (Fig. 4A, left). The correlation is not perfect and there likely are other contributing factors. Also, ligand structures are dynamic and it is difficult to quantify effective agonist volume from only a chemical formula. Nonetheless, it appears the length of the agonist’s tail and the distribution of charge in the head group are less important than head-group volume. It may be possible to predict an ligand’s efficiency by comparing its head group to others having a known efficiency. For example, adding a small group to the pyrimidine of nicotine might change affinity and efficacy but probably not efficiency.

The correlation between head-group size and efficiency aligns with results from molecular dynamics simulations of AChR binding site homology models occupied by different agonists (Fig. 4A, right) (Tripathy, Zheng, & Auerbach). In these simulations, the mean volume of the C binding pocket is similar with 51%- and 42%-efficient agonists. The pocket volume also is, in all cases, smaller in O versus C, but the extent of this reduction is greater for higher versus lower efficiency ligands. Overall, the results suggest that agonist efficiency is correlated with the extent of pocket contraction that in turn is correlated inversely with agonist head-group volume. Although these investigations need to be extended, one possible explanation is that large agonists limit pocket contraction by steric hindrance, to limit the fraction of binding energy delivered to the next steps of the gating machinery (see below).

Agonist efficiencies have been measured in AChRs having a binding site mutation (Fig. 4B). Mutations of a conserved glycine near the binding pocket lowers efficiency from 54% or 50% to 42%, the same as Ebd in wild-type (wt) receptors (Jadey, Purohit, & Auerbach, 2013). In addition, K_dC_ and K_dO_ (for ACh) have been measured following mutations of aromatic residues that line the principle side of the binding pocket (Purohit, Bruhova, Gupta, & Auerbach, 2014). Many of the substitutions have little or no effect on efficiency, but those of bulky tryptophan *α*W149 increase η to 55% (same as TMA in wt) and αY190A reduces η to 35% (the same as varenicline in wt). Perhaps mutations that change η mark the pathway for energy transfer out of the binding pocket.

It is worth noting that of the above ∼50 different binding site mutations, the resulting η-values fall into one of the 4 classes apparent in wt binding sites, namely 54, 51, 42 or 35% (Fig. 3). Thus, in addition to being discrete and precise, agonist efficiencies appear to be modal. Combining this observation with the pocket volume-change results leads to the following speculation. Whereas the shape of C pocket is mostly independent of agonist efficiency, the O pocket can adopt only a discrete number of target structures that correlate with the modal efficiency values. Perhaps the adult-type AChR pocket in the C conformation is large enough to accommodate all efficiency classes, but the contraction upon opening is frustrated by larger ligands so that different, discrete binding structures are targeted. In this sense, the distribution of efficiencies constitutes a binding site’s spectrum or ‘bar code’. Additional efficiency measurements, structures and simulations are needed to test this speculation.

### Catch-and-hold

That agonists with different affinities and efficacies have the same efficiency implies that events in binding and gating (the x- and y-axes of an efficiency plot, Fig. 3) are related energetically. It is easy to imagine the difference between these processes that are the formation of a ligand complex and the receptor isomerization, but harder to conceive of this energy link. To explore the energy linkage that is inherent to concept of agonist efficiency we need to consider events that take place inside ‘binding’ and ‘gating’.

The second unexpected set of results pertains to the kinetics of agonist binding to C versus O sites (Fig. 6). The association rate constant (k_on_) to C is below the limit set by diffusion and is agonist dependent (weaker agonists are slower to bind) even though the ligands have approximately the same size, charge, shape and diffusion constant (Nayak & Auerbach, 2017). For example, k_on_ to C for choline is ∼100 times slower than for ACh and ∼1000 times slower than the diffusion limit (Fig. 5A left). Certainly agonists must reach the binding site by diffusion, but in addition it appears that the formation of the low-affinity complex requires a local rearrangement of the binding site called ‘catch’ (Jadey & Auerbach, 2012). Another indication of a catch rearrangement is that in AChRs having the mutation αG153S, k_on_ to C for choline is highly temperature-dependent (Gupta & Auerbach, 2011). Whereas the energy change associated with the diffusional component of k_on_ is about the same for all small agonists, that for the catch rearrangement varies substantially. In GABA and glutamate receptors, too, k_on_ to C is slower than diffusion (Jones, Jonas, Sahara, & Westbrook, 2001; Robert & Howe, 2003).

In contrast, the rate constants for agonist dissociation (k_off_) from C are approximately agonist-independent. Hence, the agonist-dependence of K_dC_ (=k_off_/k_on_) is determined mainly by the energy change in catch. These results suggest that association to C involves two sequential events, diffusion and complex formation, separated by an undetected, short-lived intermediate state (Fig. 5B left). The value of logK_dC_ is the sum of agonist-dependent energy change (from catch) plus an energy offset that is the same for all agonists (from diffusion).

Agonists can also bind to and dissociate from the O conformation (Fig. 1B). Constitutive P_O_ can be increased by substituting side chains far from the neurotransmitter binding sites, for instance in the transmembrane domain. L_0_ increases if the substitutions are locally more stable in O compared to the wild type. Moreover, multiple mutations often act independently and have an additive effect on L_0_, and therefore L_2_ and EC_50_. Indeed, so far all AChR mutations away (>12 Å) from the binding sites change EC_50_ exclusively by changing L_0_ rather than the coupling constant (Gupta, Chakraborty, Vij, & Auerbach, 2017; Purohit & Auerbach, 2009).

Fig. 6A shows k_on_ and k_off_ for an O binding site in constitutively active AChRs. It was doubly-surprising to learn from direct measurements that k_on_ to O is both approximately agonist-independent and at the diffusion limit even if this was predicted earlier by using an indirect approach (Grosman & Auerbach, 2001). Apparently, these agonists indeed do simply diffuse into the O binding site. Unlike binding C, binding to O does not require a catch rearrangement. Rather, these results suggest that the distant mutations that increased L_0_ also enforced catch, to make k_on_ to O limited only by diffusion.

I now turn to gating which, like binding to C, is a multi-step process (Fig. 5B right). *A priori*, the global isomerization of the 300 kD, 5 subunit monster that is a nicotinic AChR requires passage through a multitude of intermediate transitions and short-lived intermediate states. ϕ-value analyses (see Fig. 8, below) of diliganded AChR gating indicate there are 4 major intermediate configurations between C and O (5 major transitions) that possible comprise the ∼μs gap that has been observed directly. These intermediates can be envisioned as corrugations (small bumps and dips) on the gating energy landscapes (Fig. 1A). In channel-opening, the first transition, ‘hold’, is the local rearrangement of the binding site that increases affinity, weak-to-strong. Whereas catch sets affinity, hold sets the coupling constant and, hence, efficacy. The energy changes in the remaining intermediate steps of gating involve rearrangements away from the binding sites and are agonist independent (see Fig. 8).

Only two intermediate rearrangements, one in binding and one in gating, have an agonist dependent energy change, catch and hold. These energies are proportional to (logK_dC_) and (logK_dO_-logK_dC_), with a sum (logK_dO_) that is proportional to the overall energy change for both rearrangements. Hence, a group of agonists having the same efficiency indicates that for all, the energy change in hold is the same fraction (η) of the overall energy change in catch+hold, which leads to Eq. 3.

Fig. 7A shows these energy changes as energy landscapes for a strong and a weak agonist. η is the same for agonists having different affinities and efficacies because catch and hold are linked by a linear free energy relationship (LFER). The rearrangements associated with the agonist-dependent parts of binding and gating split the agonist energy, with the fraction of the total apportioned to gating being equal to η. Strong agonists ‘tilt’ the LFER more so than weak ones, but the fractional split between catch and hold, as indicated by the position of AC in the reaction co-ordinate, remains the same for agonists having the same η.

In AChRs, structural underpinnings of the catch and hold energy changes are not known. Hold may involve an agonist-dependent contraction of the binding pocket, but other conformational changes likely also contribute. The energy relationship between hold and catch suggests that some of the structural changes in hold might also pertain to catch. To fully understand the structural basis of the energy changes in catch and hold will require knowledge of 4 configurations of the binding sites: apo-C (no agonist), A--C (with agonist but before complex formation), AC (low affinity complex) and AO (high affinity complex). A--C is of particular interest because this configuration is the agonist-dependent trigger for catch and the rest of the channel-opening process.

Fig. 7B shows progressive changes in the catch-and-hold energy landscape. The top, blue line pertains to the unliganded condition, where the catch barrier is large. The middle, grey line is the landscape after an agonist has diffused to occupy a site that is near, but outside of, the pocket. In this A--C configuration, the catch barrier is small and the formation of the low affinity AC complex occurs rapidly. The bottom, black line is after catch, which lowers the hold barrier so that the binding site switches rapidly from the low to the high affinity AC’ configuration. In brief, agonists promote catch that promotes hold.

Fig. 7B shows these events as a reaction scheme having dimensions that reflect agonist diffusion (grey), catch (blue) and hold (green), with arrows indicating the magnitude of the equilibrium constant. Because of the LFER, the predominant activation sequence after diffusion catch-then-hold. The arrows in the bottom (liganded) face indicate that hold-before-catch is rare, but also that catch-after-hold is highly favored so that the cycle is balanced. Hence, when in channel-closing a receptor finds itself in a hold-without-catch conformation, catch happens quickly. Microscopic reversibility dictates that if catch promotes hold, then hold promotes catch.

### Zipper mechanism

Efficiency shows that catch and hold rearrangements are linked by a LFER. In this section I extend this conclusion and propose that the other components within the gating isomerization, too, are linked by LFERs.

Fig. 8 shows the third surprising result, that energy changes at the AChR gating TS are organized both with regard to their value and location in the receptor. The intermediate steps of AChR gating can be inferred from the parameter ϕ determined from the slope of a log-log plot of the change in the opening rate versus gating equilibrium constant for a series of mutations of one residue. ϕ is a measure of the relative energy character of the mutated side chain at the TS as being like that of O versus C (Fig. 8A, inset). Fig. 8A shows that there are 5 discrete populations of ϕ, each being related to a contiguous set of amino acids (a domain) in the receptor. ϕ values indicate the sequence of intermediate energy changes (domain rearrangements) on a scale from 1 (early) to 0 (Fig. 8B) (Auerbach, 2005). The organized map of ϕ is a surprise and suggests that in AChR gating, 5 domains change conformation in sequence to connect the binding sites with the gate by a conformational ‘wave’ (Fig. 8C) (Gupta et al., 2017; Purohit, Gupta, Jadey, & Auerbach, 2013).

C and O structures of AChRs and related receptors suggest the nature of the specific ϕ-domain rearrangements (Calimet et al., 2013; Gharpure, Noviello, & Hibbs, 2020; Taly et al., 2005; Yu et al., 2021). In opening, the first is ‘hold’ that likely involves a contraction of the binding pocket. This is followed by ‘twist’ (a rotation and compaction of the extracellular domain), ‘tilt’ (a rearrangement of transmembrane helices), ‘rip’ (a change in the hydrophobicity of the gate region), and ‘pop’ (fusion of the extra- and intracellular water compartments to initiate ion conduction). In channel closing, these rearrangements occur in the reverse order.

Together, the 5 domain transitions comprise a conformational cascade. The ϕ distribution and map divides the overall gating TS into a series of intermediate transitions in specific structural domains but does not offer lifetimes for the 4 intermediate, high-affinity pre-open states. If the ∼μs shut interval indeed reflects sojourns within the gating cascade, then the intermediate states could have lifetimes of ∼100 ns.

Given that catch and hold are linked by a LFER, it reasonable to speculate that the components of the gating cascade, too, are linked energetically. The speculation is that in the binding-gating activation sequence catch-hold-twist-tilt-rip-pop, each pair of steps is linked by a LFER so, like catch-and-hold, each transition promotes its neighbor. In this view, everything between diffusion and water fusion is linked energetically.

In a zipper, the barrier to the formation of the next pair of interdigitating teeth is lowered by the formation of the adjacent, preceding pair. In the above model the entire chain of binding and gating rearrangements is linked similarly. In receptor activation, the agonist lowers the catch local barrier (perhaps by structure matching), which lowers the hold barrier, which lowers twist, then tilt and rip, which finally lowers the barrier to joining the water compartments. Just as in a zipper, each linkage element lowers the barrier to forming the next in the chain, with zippering able to be completed, C to O, in ∼μs. Because of microscopic reversibility, the predominant pathway for receptor deactivation is the reverse process, pop-rip-tilt-twist-hold-catch.

Any proposal regarding the mechanism of AChR activation must account for the facts that i) there is correlation between affinity and efficacy (Fig. 3) and ii) there are 5 ϕ populations that decrease domain-wise, binding site to gate (Fig. 8). The zipper mechanism not only accounts for these but also the observation that k_on_ to O is diffusion-limited. A mutation that stabilizes O, for example by promoting water fusion at the gate, will stabilize all of the other elements in the chain, all the way back to catch, across the binding gating divide.

Indeed, with regard to energy there is no such divide. If each rearrangement occurred independently of the others, efficiency would not exist, activation would be slow and the distribution of ϕ would be disorganized rather than modal. However, because of the LFERs and microscopic reversibility the model predicts a simple forward and backward march through a linear reaction chain that generates the affinity-efficacy correlation and an organized map of ϕ.

Efficiency is the experimental manifestation of the fractional conversion of binding energy from catch to hold. Presumably each link in the zipper, too, has an energy conversion efficiency. What makes agonist η remarkable is that the components of this LFER belong to “binding” and “gating”, processes that have long been separated in our conception of receptor activation. Efficiency reveals that these processes are stages of a single, energy-linked cascade of domain rearrangements. In terms of energy (structure), binding and gating are parts of a single, reversible conformational sweep, agonists to ions.

Certainly, all of these energy linkages need to be identified as structure interactions. Where does the agonist bind to trigger catch? What is the catch rearrangement? What are the structural correlates of an agonist lowering the barrier to catch, and catch to hold? Similar questions apply to the rest of the ϕ-domain transitions that populate the activation pathway. What structural interactions determine that each step promotes the rearrangement of the next element in the chain? Structures reveal stable states, kinetics reveals transition states and energies (calculated from structures and kinetics) reveal the forces that drive movements. It will take time to isolate, identify, solve and understand the intermediates of the AChR zipper.

